## Supplementary figures 1-6, Supplementary Table 1, Figure Legend for Supplementary Table 2 and Supplementary Document 2 for "Behavioral screening defines three molecular Parkinsonism subgroups in *Drosophila*"

Suppl Fig 1

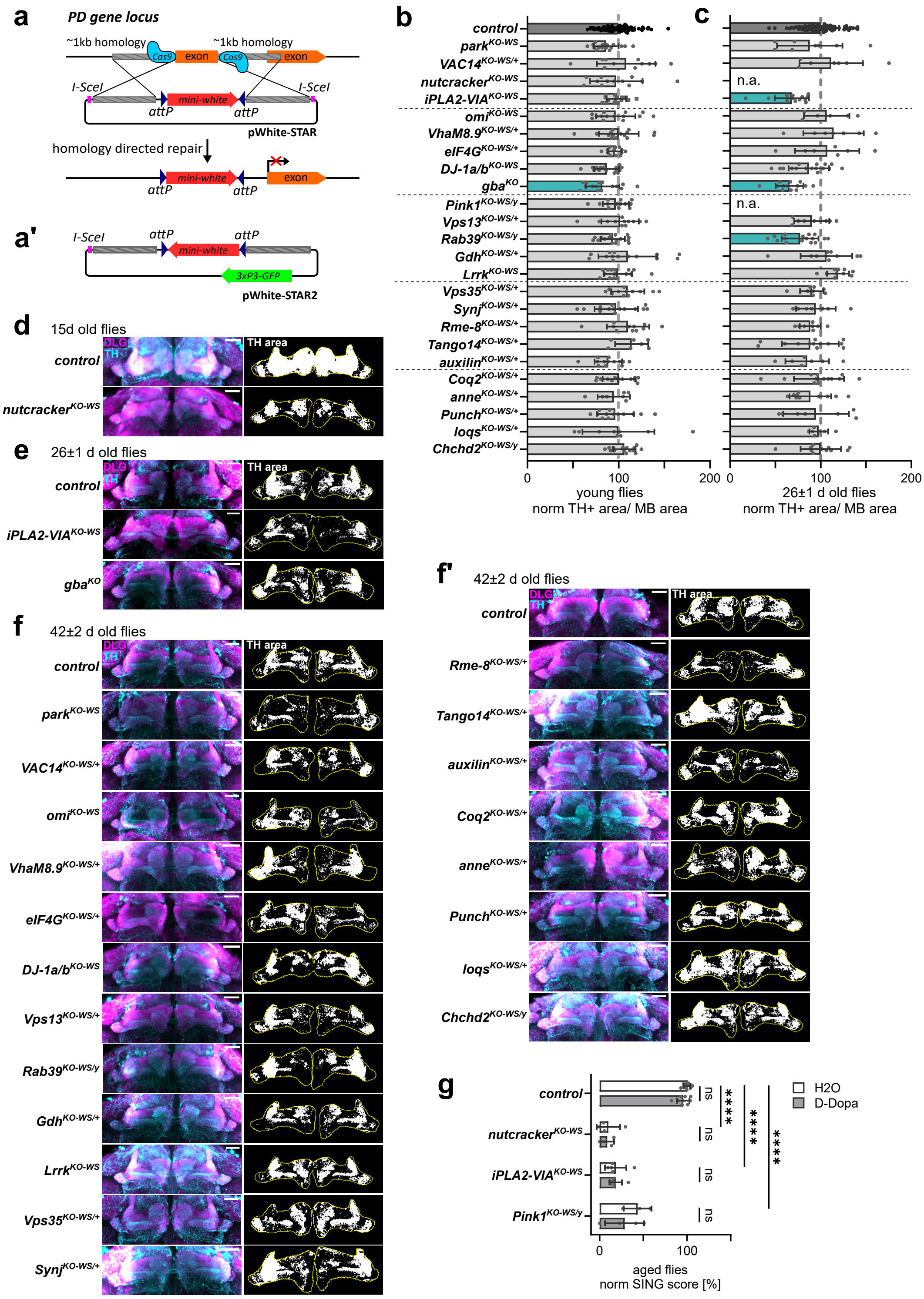


**Supplementary Fig.1: Parkinsonism fly collection shows age-dependent dopaminergic defects**, *related to Fig.1*. (a-a') Scheme of the genetic approach for the generation of the parkinsonism mutant fly collection. The first common exon of all possible *Drosophila* transcripts of the targeted parkinsonism gene was replaced by attP-flanked mini-*white* gene, by homology directed repair with the pWhite-STAR (a) or pWhite-STAR2 (a') using CRISPR/Cas9, creating a null mutant. I-SceI sites were utilized in the rare event of full donor plasmid integration (methods). (b,c) Quantification of dopaminergic synaptic area within MB area at (b) 5±1 d and (c) 26±1 d after eclosion in mutants relative to the control. Due to their shorter lifespan *nutcracker*$^{KO\text{-}WS}$ were tested at 15 d, *Pink1*$^{KO\text{-}WS}$ at 22±2 d and are not included in (c). Turquoise colored bars represent $p<0.05$, (b) One-way ANOVA with Dunnett's test and (c) ANOVA Kruskal-Wallis with Benjamini-Hochberg. Bars: mean ± SD; points are individual animals N≥6 per genotype. (d-f) Maximum projection confocal images of control and mutant fly brains at (d) 15±1 d, (e) 26±1 d and (f-f') 42±2 d after eclosion stained with anti-TH (cyan) and anti-DLG (magenta) antibodies, where DLG marks the post-synaptic site of MB. The black and white images represent the thresholded TH area (in white) of "middle z-plane" within the ROI (yellow, outline of MB). Scale bar: 20 µm. (g) SING quantification of aged flies treated with solvent control H2O or D-Dopa 10 d prior to the assay relative to $H_2O$ treated control. Bars: mean ± SD; points represent groups of animals and N≥3. Two-way ANOVA with Tukey's multiple comparison: ns, not significant; **** $p<0.0001$. DLG, Discs-large; MB, mushroom body; SING, startle-induced negative geotaxis; TH, tyrosine hydroxylase.

Suppl Fig 2

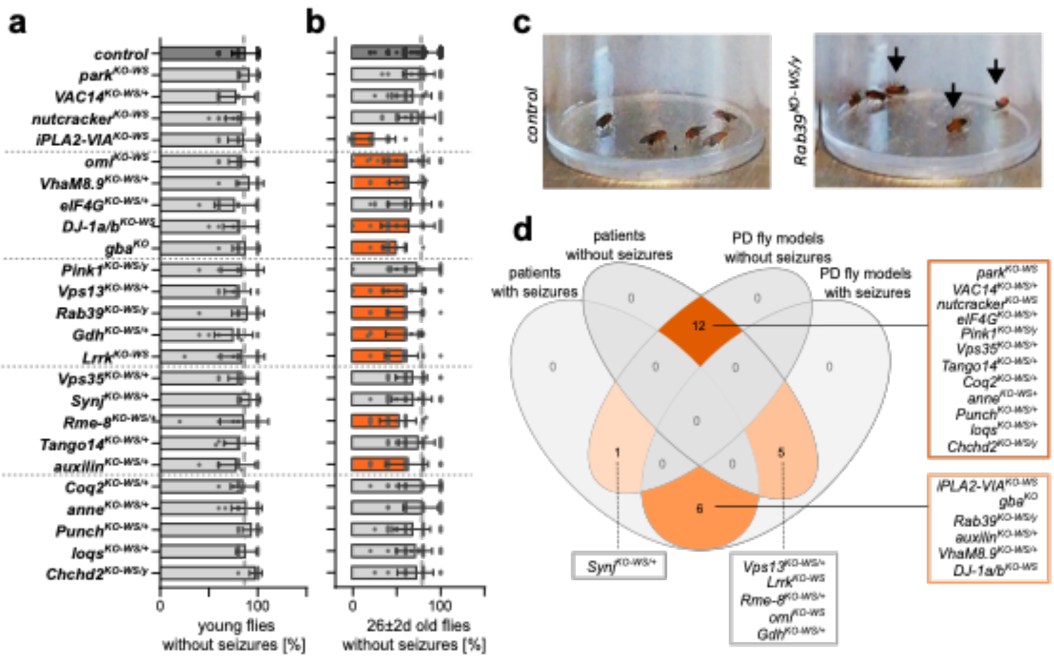


**Supplementary Fig.2: Several parkinsonism mutants show seizure-like behavior**, *related to Fig.1.* (a,b) Quantification of parkinsonism mutants without seizure-like behavior after sensory stimulation (vortex) at (a) 6±1 d (young) and (b) 26 ±2 d after eclosion (except for *nutcracker*$^{KO\text{-}WS}$ at 15±1 d and, *Pink1*$^{KO\text{-}WS}$ at 22±2 d). Orange colored bars represent $p \leq 0.01$, ANOVA Kruskal-Wallis with Benjamini-Hochberg compared. Bars: mean ± SD; points are groups of animals and $N \geq 9$. (c) Representative images of 25±1 d old control and *Rab39*$^{KO\text{-}WS}$ after seizure induction, black arrows indicate flies with seizure-like phenotypes. (d) Venn diagram representing the convergence of patients with familial forms of parkinsonism described to suffer from seizures and fly mutants with or without seizure-like behavior.

Suppl Fig 3

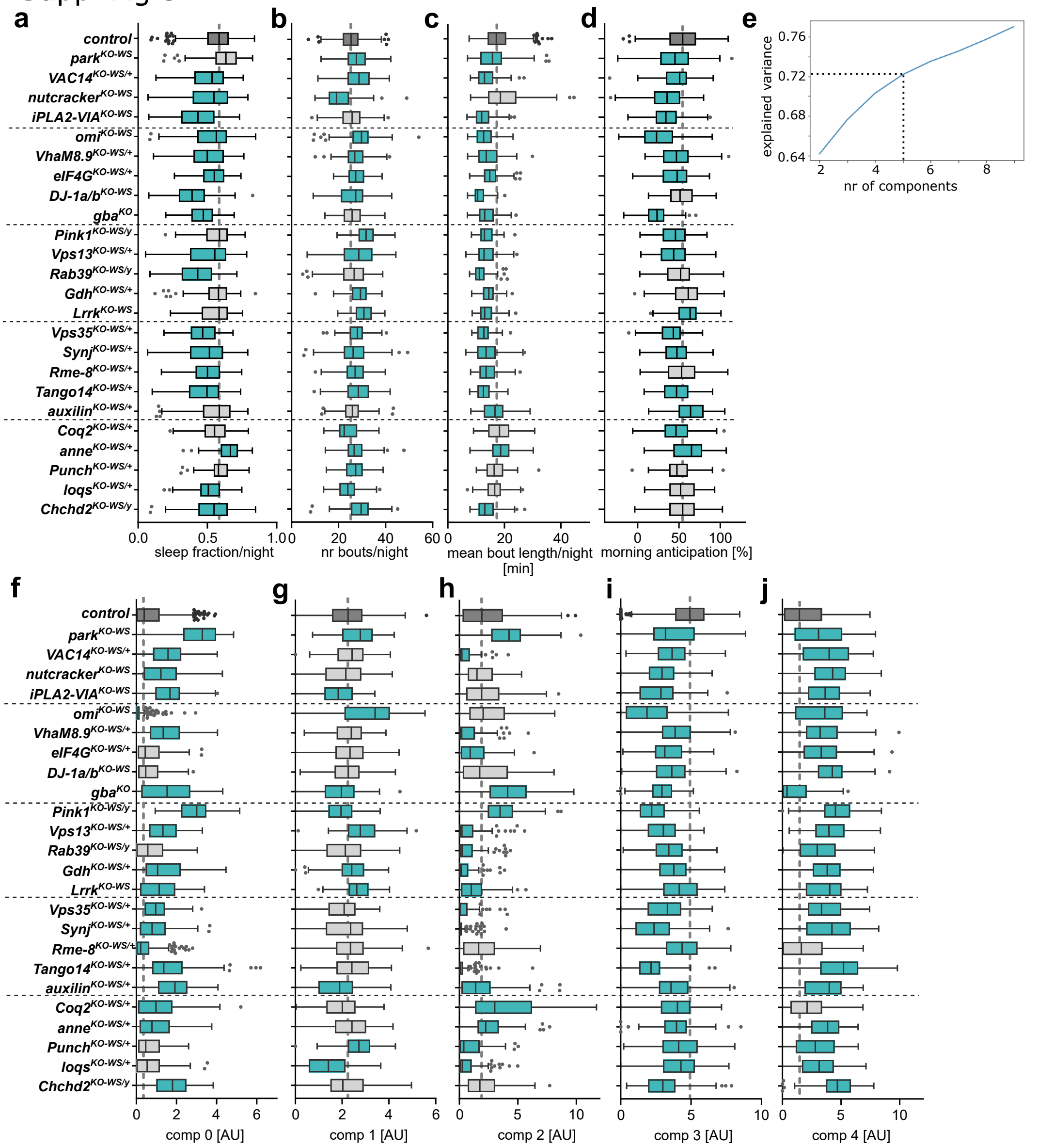


**Supplementary Fig.3: Young parkinsonism mutant flies display diverse behavioral defects**, *related to Fig.2*. Quantification of (a) sleep fraction per night, (b) number of bouts per night, (c) mean bout length per night in min and (d) morning anticipation in % in parkinsonism mutants averaged over 5 d. (e) Elbow method supports 5 components as meaningful number of features explaining 72 % of the variance. (f-j) Quantification of components 0-4 represented by single flies in individual genotypes. (a-d, f-j) Box and whisker plot (Tukey method): median and IQR; statistical significance: ANOVA Kruskal-Wallis with Benjamini-Hochberg; turquoise colored boxes represent $p<0.05$, from individual animals $N\geq80$.

Suppl Figure 4

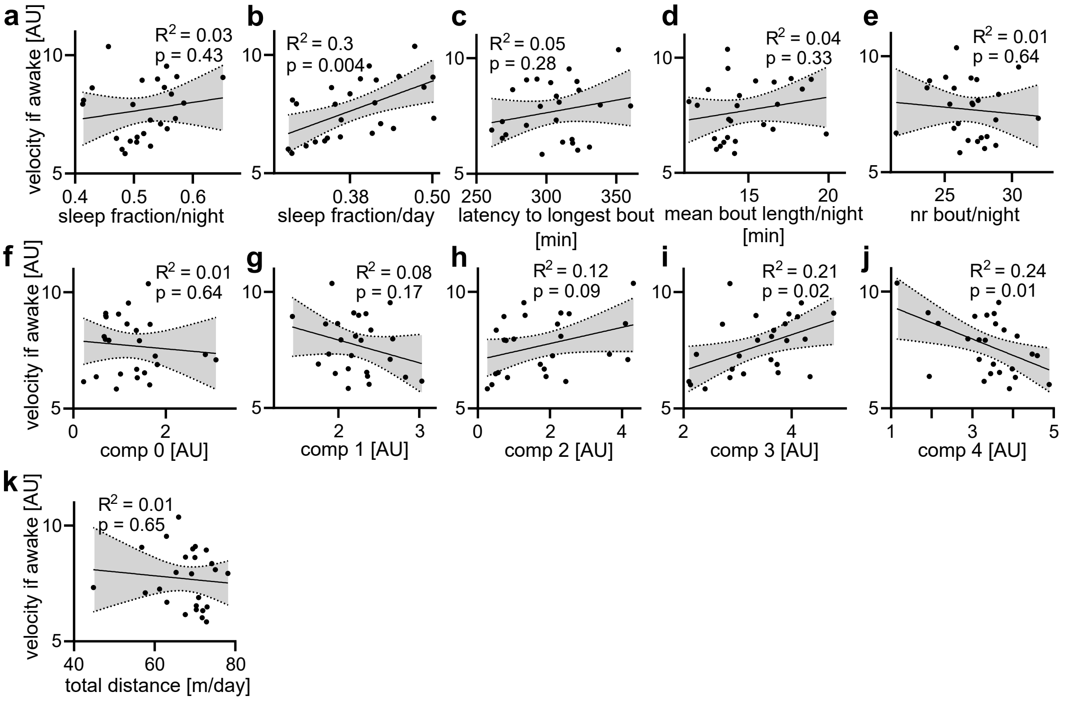


**Supplementary Fig.4: Basic motor performance does not generally correlate with behavioral parameters or NMF components 0-4**, *related to Fig.2*. Linear regression plots with 95% confidence intervals (grey area) of behavior parameters with velocity if awake [AU] (a-k). Dots represent the mean of parkinsonism mutants. $R^2$ and p-values are indicated.

Suppl Fig 5

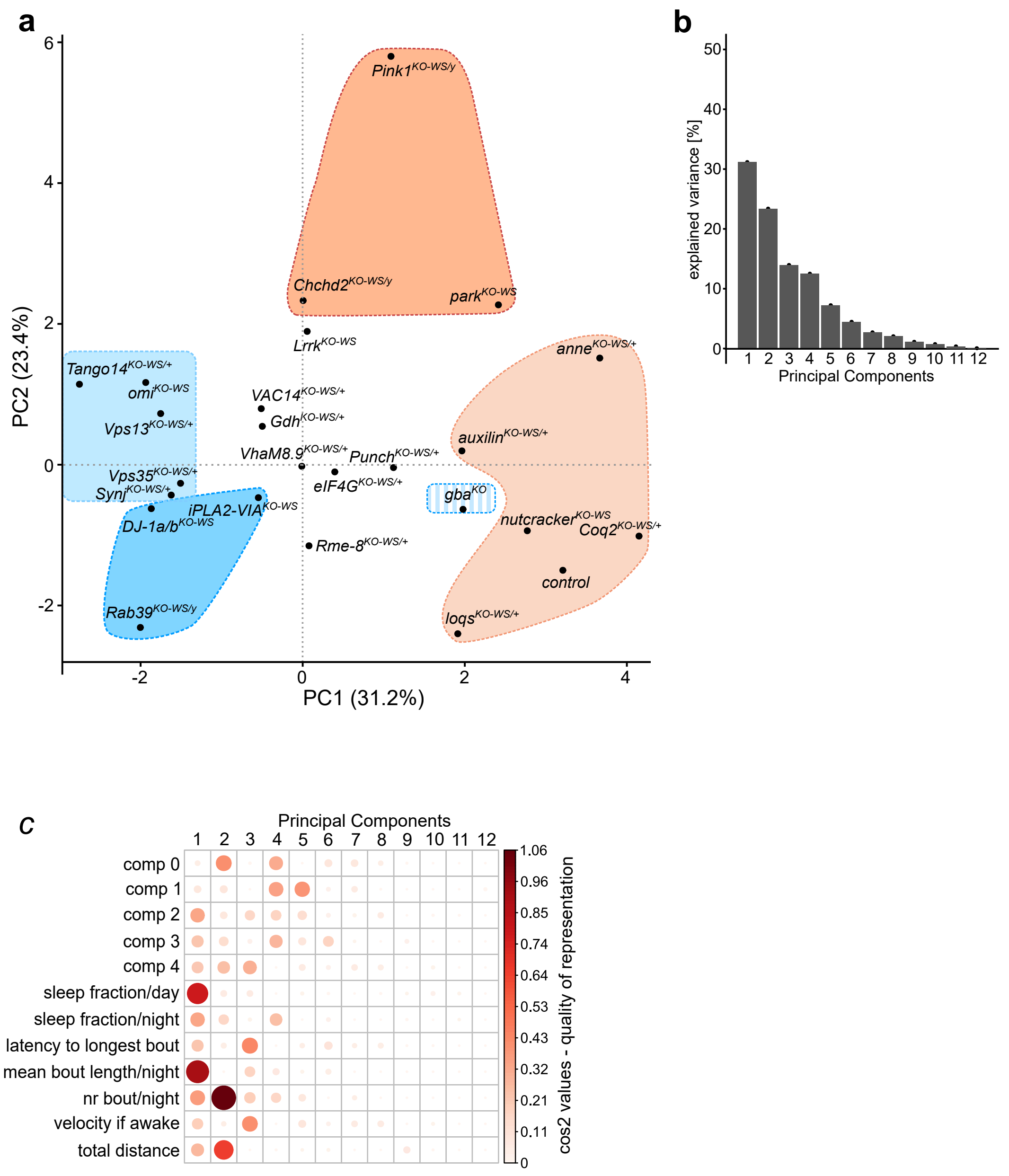


**Supplementary Fig.5: Principal component analysis (PCA) supports the assignment of genes to groups identified with hierarchical clustering**, *related to Fig.2*. (a) PCA plot of parkinsonism mutants based on their scaled behavior features and boxes are inserted manually highlighting behavior groups, color coded as in Fig.2d. Principal component (PC) 1 and 2 explain 55% of total variance. (b) Screen plot describing the explained variance of individual PCs. (c) Cos2 plot describing the quality of representation of behavior parameters within individual PCs. Both dot size and color indicate the cos2 value. Comp, component; nr, number.

Suppl Fig 6

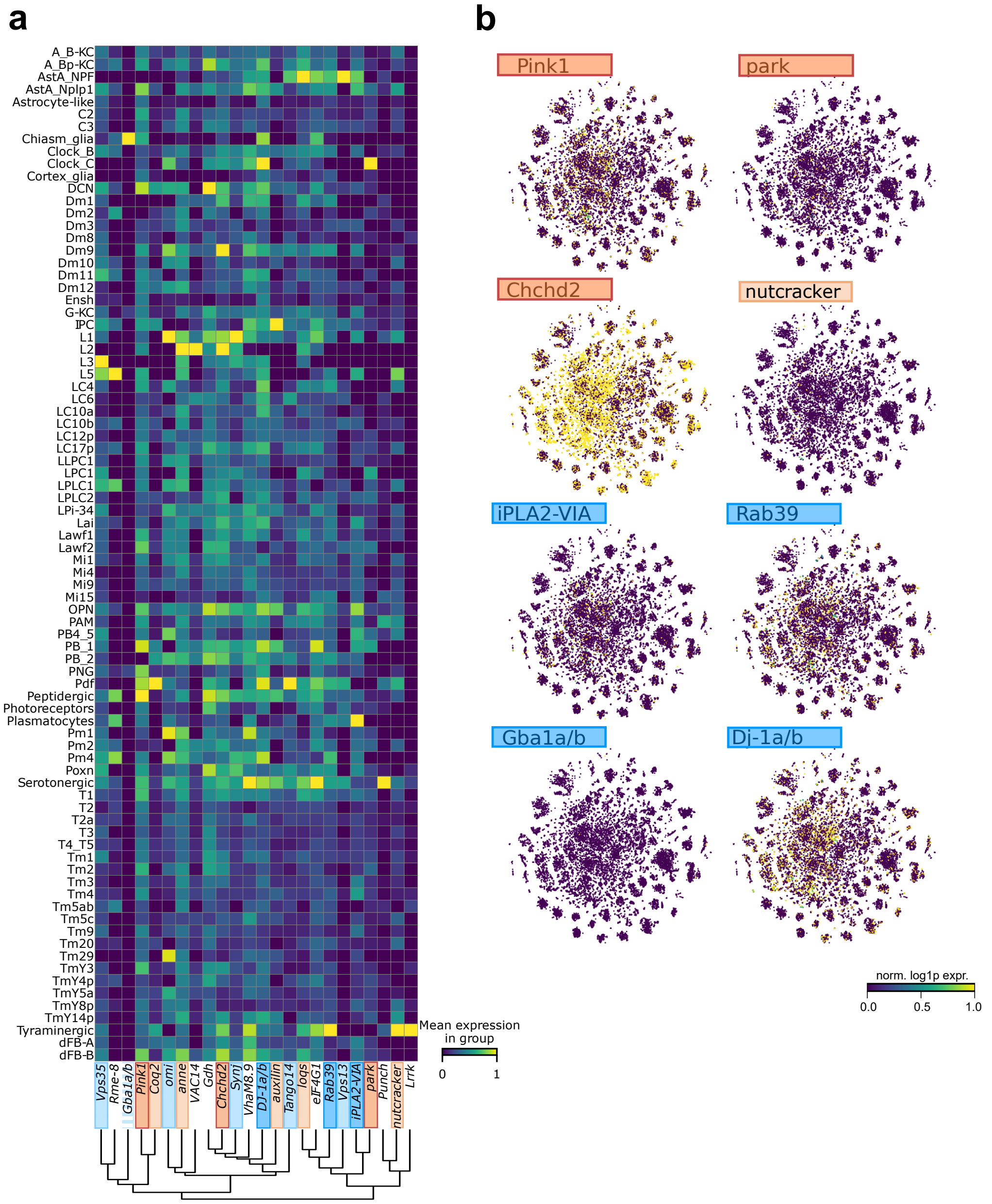


**Supplementary Fig.6: Parkinsonism gene expression clustering across cell-types of the fly brain.** (a) Heatmap and hierarchical clustering of scaled parkinsonism gene expression across all identified cell types in single-cell RNAseq atlas[28] of young (5±1 d) control flies. Parkinsonism genes are color coded according to behavior groups. (b) Example UMAPs of *Pink1*, *park*, *Chchd2*, *nutcracker*, *iPLA2-VIA*, *Rab39*, *gba1a/b* and *DJ-1a/b* normalized and logarithmized expression values.

**Supplementary Table 1: Overview of genes with mutations in familial parkinsonism conserved in *D. melanogaster*,** *related to Fig.1*.

*Nomenclator in paper, Typical: bradykinesia, resting tremor, rigidity, postural instability

| hGene name | dGene name | inheritance | OMIM | disease onset | motor phenotype | additional features |
|---|---|---|---|---|---|---|
| ***ATP13A2*** *(PARK9)* | *anne* | AR | 606693 | juvenile | atypical | Kufor-Rakeb syndrome, dementia, spasticity |
| ***ATP6AP2*** | *VhaM8.9* | XLR | 300911 | variable | typical | Spasticity, seizures |
| ***CHCHD2*** *(PARK22)* | *Chchd2* | AD | 616710 | adult | typical | |
| ***COQ2*** | *Coq2* | AD, AR | 146500 | adult | typical | MSA susceptibility |
| ***DJ-1*** *(PARK7)* | *DJ-1a* | AR | 606324 | early | typical | seizures |
| | *DJ-1b* | | | | | |
| ***DNAJC13*** *(PARK21)* | *Rme-8* | AD | 616361 | adult | typical | |
| ***DNAJC6*** *(PARK19A/B)* | *auxilin* | AR | 615528 | A: juvenile<br>B: early | typical | A: mental retardation and seizures<br>B: typical PD features |
| ***eIF4G1*** *(PARK18)* | *eIF4G* | AD | 614251 | adult | typical | |
| ***HTRA2*** *(PARK13)* | *omi* | AD | 610297 | adult | typical | |
| ***LRRK2*** *(PARK8)* | *Lrrk* | AD | 607060 | adult | typical | |
| ***parkin*** *(PARK2)* | *park* | AR | 600116 | juvenile | typical | |
| ***PINK1*** *(PARK6)* | *Pink1* | AR | 605909 | early | typical | |
| ***PLA2G6*** *(PARK14)* | *iPLA2-VIA* | AR | 612953 | early | atypical | dystonia-parkinsonism with extrapyramidal features, psychiatric/cognitive issues, possible brain iron accumulation, seizures |
| ***Rab39B*** | *Rab39* | XLR | 311510 | early | atypical | Waisman syndrome: mental retardation, seizures |
| ***Synj1*** *(PARK20)* | *Synj* | AR | 615530 | early | typical | seizures, cognitive decline, dystonia |
| ***VPS35*** *(PARK17)* | *Vps35* | AD | 614203 | adult | typical | |
| ***PRKRA*** | *loqs* | AR | 612067 | early | atypical | dystonia-parkinsonism |
| ***FBXO7*** *(PARK15)* | *nutcracker* | AR | 260300 | early | atypical | parkinsonian-pyramidal syndrome, spasticity, hyperreflexia |
| ***VAC14*** | *CG5608 (VAC14*)* | AR | 617054 | juvenile | atypical | Striatonigral degeneration with dystonia |
| ***VPS13C*** *(PARK23)* | *Vps13* | AR | 616840 | early | atypical | cognitive decline, visual hallucinations |
| ***Gch1*** | *Punch* | AD, AR | 128230 | juvenile | typical | DOPA-responsive dystonia |
| ***GLUD2*** | *Gdh* | Risk gene | 300144 | early | typical | |
| ***GBA*** | *gba 1a & 1b (gba*)* | Risk gene | 168600 | adult | typical | Gaucher disease; susceptibility to PD with cognitive decline, visual hallucinations and psychiatric symptoms, seizures |
| ***NUS1*** | *Tango14* | Risk gene | Guo et al. 2018 | early | typical | |

**Supplementary Table 2: Average Genetic Interaction strength of all gene pairs including Bayesian estimate of interaction significance**, *related to Fig.3*.

"Expected" relates to the expected normalized depolarization according to the non-interacting model based on single heterozygous mutants, while the "mean GI strength" is the subtraction of the modelled expected depolarization from the individual observed depolarization amplitude of heterozygous gene pairs. Hdi0 and hdi1 delineate the limits of the 95% high-density interval (HDI). Dist_0 show the distance from the HDI to zero. If dist_0 = 0 the interval contains zero. The gene pair $Pink1^{KO\text{-}WS/y}$ and $park^{KO\text{-}WS/+}$ showed synthetic lethality and was thus scored -2.

**Supplementary Document 2: Cloning and primer details for parkinsonism KO *Drosophila* models,** *related to Method details*

**Cloning of *pWhite-STAR2***

*Primers to generate SV40 poly(A)-tail*

F_hifi_SV40: GGTGGTCCCGTCGAAAGCCGAGGATCCAGACATGATAAG

R_hifi_SV40: ATGGACGAGCTGTACAAGTAAGGATCTTTGTGAAGGAAC

gBlock_P3XGFP:
CGGCCGCTATAACTAGTAACTCGAGGGATCTAATTCAATTAGAGACTAATTCAATTAGAG
CTAATTCAATTAGGATCACAGCTTATCGATTTCGAACCCTCGACCGCCGGAGTATAAATA
GAGGCGCTTCGTCTACGGAGCGACAATTCAATTCAAACAAGCAAAGTGAACACGTCGCT
AAGCGAAAGCTAAGCAAATAAACAAGCGCAGCTGAACAAGCTAAACAATCGGCAAAAT
GGTGAGCAAGGGCGAGGAGCTGTTCACCGGGGTGGTGCCCATCCTGGTCGAGCTGGACG
GCGACGTAAACGGCCACAAGTTCAGCGTGTCCGGCGAGGGCGAGGGCGATGCCACCTAC
GGCAAGCTGACCCTGAAGTTCATCTGCACCACCGGCAAGCTGCCCGTGCCCTGGCCCACC
CTCGTGACCACCCTGACCTACGGCGTGCAGTGCTTCAGCCGCTACCCCGACCACATGAAG
CAGCACGACTTCTTCAAGTCCGCCATGCCCGAAGGCTACGTCCAGGAGCGCACCATCTTC
TTCAAGGACGACGGCAACTACAAGACCCGCGCCGAGGTGAAGTTCGAGGGCGACACCCT
GGTGAACCGCATCGAGCTGAAGGGCATCGACTTCAAGGAGGACGGCAACATCCTGGGGC
ACAAGCTGGAGTACAACTACAACAGCCACAACGTCTATATCATGGCCGACAAGCAGAAG
AACGGCATCAAGGTGAACTTCAAGATCCGCCACAACATCGAGGACGGCAGCGTGCAGCT
CGCCGACCACTACCAGCAGAACACCCCCATCGGCGACGGCCCCGTGCTGCTGCCCGACAA
CCACTACCTGAGCACCCAGTCCGCCCTGAGCAAAGACCCCAACGAGAAGCGCGATCACA
TGGTCCTGCTGGAGTTCGTGACCGCCGCCGGGATCACTCTCGGCATGGACGAGCTGTACA
AGTAA

**Cloning of pReC-CG31413**

*gBlocks to generate pReC*

gBlock_attB-MCS-SV40:
GTTGTAAAACGACGGCCAGTGAATTCCCGCGGTGCGGGTGCCAGGGCGTGCCCTTGGGCT
CCCCGGGCGCGTACTCCACTCGAGCGCTAGCTAATCGTCTAGatgcGAAGAGCGAGCTCGG
TAGATCCGATATCCTGCAGGCATGCAAGCTTGGCGTAATCATGGCTCTTCgtaaGGATCTTT
GTGAAGGAACCTTACTTCTGTGGTGTGACATAATTGGACAAACTACCTACAGAGATTTAA
AGCTCTAAGGTAAATATAAAATTTTTAAGTGTATAATGTGTTAAACTACTGATTCTAATTG
TTTGTGTATTTTAGATTCCAACCTATGGAACTGATGAATGGGAGCAGTGGTGGAATGCCT
TTAATGAGGAAAACCTGTTTTGCTCAGAAGAAATGCCATCTAGTGATGATGAGGCTACTG
CTGACTCTCAACATTCTACTCCTCCAAAAAAGAAGAGAAAGGTAGAAGACCCCAAGGAC
TTTCCTTCAGAATTGCTAAGTTTTTTGAGTCATGCTGTGTTTAGTAATAGAACTCTTGCTTG
C

gBlock_SV40-attB:
AGTAATAGAACTCTTGCTTGCTTTGCTATTTACACCACAAAGGAAAAAGCTGCACTGCTA
TACAAGAAAATTATGGAAAAATATTTGATGTATAGTGCCTTGACTAGAGATCATAATCAG

CCATACCACATTTGTAGAGGTTTTACTTGCTTTAAAAAACCTCCCACACCTCCCCCTGAAC
CTGAAACATAAAATGAATGCAATTGTTGTTGTTAACTTGTTTATTGCAGCTTATAATGGTT
ACAAATAAAGCAATAGCATCACAAATTTCACAAATAAAGCATTTTTTTCACTGCATTCTA
GTTGTGGTTTGTCCAAACTCATCAATGTATCTTATCATGTCTGGATCCGTGGAGTACGCGC
CCGGGGAGCCCAAGGGCACGCCCTGGCACCCGCACCGCGGGGTACCGTCATAGCTGTTTC
CTGTGTGAAATTGTTATCCGCTCACAATTCCACACAACATACGAGCCGGAAGCATAAAGT
GTAAAGCCTGGGGTGCCTAATGAGTGAGCTAACTCACATTAATTGCGTTGCGCTCACTGC
CCGCTTTCCAGTCGGGAAACCTGTCGTGCCAGCTGCATTAATGAATCGGCCAACGCGCGG
GGAGAGGCGGTTTGCGTATTGGGCGCACATCCGCTTCCTCGCTCACTGACTCGCTG

*Primers to amplify the CG31413 genomic region*

F_hifi_CG31413: CTCCCCGGGCGCGTACTCCACAGAAGATGGTTGGTTTCGG

R_hifi_CG31413: TACCGAGCTCGCTCTTCgcatCTCGGGCCAAAAACAATGC

***gRNA's used to target genes of interest and make the pCFD4 constructs***

| ***Gene*** | ***Construct*** | ***5'-3' gRNA 1*** | ***5'-3' gRNA 2*** |
|---|---|---|---|
| *ATP13A2* | pCFD4-Td_gRNA:ATP13A2 | ACAATATCAAGTTTGCGCCT | CTACTTTTGAATTGGGCCG |
| *Vham8.9* | pCFD4-Td_gRNA: Vham8.9 | GTTATCGAATGGGTATCAGT | GCATTTTACTTGATTATAGA |
| *CHCHD2* | pCFD4-Td_gRNA:CHCHD2 | GTTTTGCCAGTCACAACGGG | AACATGAATGTCTCGAACGC |
| *Coq2* | pCFD4-Td_gRNA:Coq2 | ATGCGTCGTGGGTTTCTCTG | TTCTGTAAGGTAATATAACA |
| *Dj1-a* | pCFD4-Td_gRNA: Dj1-a | CGTTTTTGTTTTACAAGGGA | TTGTGCCGCTCCCACGGCGC |
| *Dj1-b* | pCFD4-Td_gRNA: Dj1-b | GCTTTGAGTTTTCCTTATCG | GCCGGACACCTCGCTGGCCC |
| *rme-8* | pCFD4-Td_gRNA: rme-8 | AGACCAGTCATGAATTGCCG | TTTAGTTCCGAGCTCTCCGT |
| *auxilin* | pCFD4-Td_gRNA: Auxilin | GCTCTTATCGATGTGGCGCG | TCACGGTTACGCGTTACCGC |
| *eIF4G1* | pCFD4-Td_gRNA: eIF4G1 | TTGATATACAGCTACCAGAG | GCAGCATAGCCATCCCACTG |
| *ntc* | pCFD4-Td_gRNA: ntc | ATATATTTGATAAGACACT | AAACTATCCTCAAAGATAA |
| *Gba1a_1b* | pCFD4-Td_gRNA: Gba1a_1b | AAACCAACCATCTTCTGGT | ATTTGAGGAGTCTCATGGCC |
| *punch* | pCFD4-Td_gRNA:punch | TAGTTAGTCAATGAACTAGG | CCGGCGTCTTGATCAGTCCC |
| *Gdh* | pCFD4-Td_gRNA:Gdh | AGTCGACTGATCAAGCCGGT | ATGAAAGTGCATCATCCTAA |
| *Omi* | pCFD4-Td_gRNA:Omi | GTATTGCTGTCGTTTCTAGT | TCCCGCCACCATCGAGGACG |
| *Lrrk* | pCFD4-Td_gRNA:Lrrk | CAGTGCGACATAAAACGGCC | GCTCTCGTAACCAGACTGGC |
| *Tango14* | pCFD4-Td_gRNA:Tango14 | TAGCAACGTAATCGATTCC | CGTAGGCACATCGTTCGCG |
| *Park* | pCFD4-Td_gRNA:Park | GCAGATCTTGCGATGGGTGC | ATTTCTGGCGCTGCACGGG |

| | | | |
|---|---|---|---|
| *Pink1* | pCFD4-Td_gRNA: Pink1 | TGCGGGGCGAAGTG TGGGGG | ATTAAACGGTGATC CCGACG |
| *iPLA2-VIA* | pCFD4-Td_gRNA: iPLA2-VIA | TTGCAAGCCCCTAC CATTGG | CCTTGGCTGATAAT CCCTCC |
| *Loqs* | pCFD4-Td_gRNA: Loqs | AACAAACTTACGAT CATGCC | TCCGCTGGTCCGTC GGTGAC |
| *Rab39* | pCFD4-Td_gRNA: Rab39 | CTACTATCGATTACT TTTGC | CTGCTCAAATTCTTC ACAGA |
| *Synj* | pCFD4-Td_gRNA:Synj | GTACTTCGTTTTTAA GGTGT | AGCTCTCTCAACAC AGTGGT |
| *VPS35* | pCFD4-Td_gRNA:VPS35 | GTATGGTGGTGACT TCTCTG | GAAGTTGTATAAAG ACGCTT |
| *VAC14* | pCFD4-Td_gRNA:VAC14 | ATTAGGATGACGAA TCTCGG | TATTTGTGACCATTA GTCAT |

**Cloning of the donor constructs:**

The homology arm gBlocks undergo modifications to ensure compatibility with the manufacturing prerequisites. Within these modified gBlocks, specific mutations are introduced, highlighted in green, strategically designed to prevent Cas9-mediated cleavage within the donor plasmid.

***Gene: ATP13A2, Constuct: pWhite-STAR_ATP13A2***

gBlock_ATP13A2_LHA:
AACAGGGTAATGGTACCTACGTTCCGCAAGAATATCTGCGGTTTCAGGTATTTTTAATAG
CTTCAAAAACCAACAAGCGTCCGATGGTGCCGTATTGTTCGCGTCTAGTCACATTAATAA
TACCCGTTACTCGTAGATTAAAAGGCTACACTAGATTTGTTGAAAAGTTTGTAACATGTA
GAAGGAAGCGTTTCCAACCATATAAAGTATATATTCTATATACATACATACATATGTATA
TTCATGTTTTTATTAATTTTCGCGTCGACTTTAGTCTGCCAGCTGAATGCTTGGCGAAAAG
TTTTAAGTAGGTTTTTAAGGCAGTTAAAGGATGAATAATGATTTAAGACATGGCGTTTAA
ATTTGATTTGTTGGCTTCGAGTAAATTGCTTTTCTATATATCGAAAATTCTCAGATATGAT
TACTGACTGCATTTTAAATTACCTACTAATTAAAAGTTTAAATAGGTGACTAATGTATGTT
CATATTTTTAAATAAATATGTATATGCATTTGTATGTATATGAAGCAACAAATCTGTTTTC
AGCGTTGTTGTATTAATATATGCATACATACATGTATTGATCAGCACAAGCCAGCTGCAC
AAAAATTTGAATGAATATACATATGTACATATGTACGGTACATGTGTATGCATTAATTGA
TTGTTTCTCATTGTATTTGTTATGACCATCTCGAATGTGTGTATAAGTGGCTCAAAATTCA
TGTATTATGTTTTCACTTTGTATGTACATATGTATGTATGGAAAGCTTCATATATGAGTGA
ATTATTAGGTGTAACGCACAATATTTCTATTAAAGATTTGTTCTGTTGGCAACAGTGTCCC
AATTACGTACAATTGAAAAGCCCCATAATCATATAAGCGAATAAAAAACGGGTACGAAA
AGAAAAGTGTGAGTGCTCTAGCGAACGAACAATATCAAGTTCGCGCTTTGGATTCTTATC
ATTTGTCTTTTATCTATTGATGTTGCCTTTTCGATTAGGCGTTTAAATAGCTGTTCGTGGCA
ACAACTTTTTCTTTAAATGTGCCTTGATTTTTTTGGTAAAAGGTCACAGTTAATTGCCTTA
ACAATGTTAGTTTCAAGACATTAACGGATTAAATTTTAGCTATGCAATTTAACAATATTTT
TTGTTTCAACTATAGGAGTGTAATTGATTTAACTTTACTGTTCTTATAATTATACAAAATA
AAACAGACAAAAATAAGCCAAGGTAGGCGTACTTAAATACATACATAGATAGAAGAAAA
TAGCAAATTATGACACCCCACAATGATTCAAAAGGTATGCACTCGATATGCGGCCGCGTA
GTGC
gBlock_ATP13A2_RHA:
CGGCCGCTATAACTAGTAACCATTCATCAGGACACTTTTGGAAAAGGAACAGCAATCCAT
AGAAAGAACTCATATAGAATGCGATCATGTAGAAAACGTCCTTCAACTGTCAGTTCATTT
TACAAGTGCCCAATTCAAAAGTACGTGCCATATAAACATATTAAGATACACCAATTTCCA
ACATCGTTTTTTTTTTAGAATGCTCATCAATACGAATTTTTCGCTGCAAGCAATTGGTTTAT
GCTTGGAACAATAATACAAATAGATTTCAAAGGATAAATGGACTCGACCTAAATATTCCT
TGTTCATATTATCACCAACAGCGTGGATTACCTGTACATGAACAGATTTCAAGGCGAATT
GTTTTCGGAGATAATGAGATAACTGTACCATTGCGAGATTTCAAGACATTGCTGTTCTTAG
AAGTACTTAATCCTTTTTACGTTTTTCAATTATTTTCTGTAATTCTTTGGTTTACATATGAT
TACTATTACTATGCTTGCGTAATACTCTTGATGTCAGTTTTTGGTATAACAGTGTCTGTTTT
ACAAACGAAAAAGGTAAGTATATCAATTAGAAGAACTTTATAAAATATAATTCCTATATA
TTTTTAGAATCAGGATGTGCTCCAAAAAACAGTATATAACACTGGTAATGCTTGGGTTGT
TGATCATAAAGGACTGTCTAAAGAGCTTCCAACGCGAGCGATAGTACCTGGGGACATCAT
TGAAATACCCTCATCAGGGTGTACGCTGCATTGCGATGCAATCTTAATATCAGGAAACTG
CATTCTAGATGAGTCTATGCTTACTGGTGAAAGTGTGCCAGTGACCAAAACTCCTCTACC
GTCGAAACGTGACATGATTTTTGATAAAACAGAGCATGCCAGACATACACTTTTTTGTGG
CACAAAGGTTATTCAGACTCGTTATATTGGCTCCAAAAAAGTATTAGCATTTGTAATAAA

CACTGGAAACATAACGGCAAAAGGAGAACTTATACGTTCTATTCTTTATCCtCCCCCTGTG
GACTACAAGTTTGAACAAGATTCGTACAAATTTATCCAGTTTCTGGCCATAATAGCATGT
GTAGGATTTATTTATACGCGTAGGGATAACAGGGTAATG

***Gene: Vham8.9, Construct: pWhite-STAR_Vham8.9***
LHA_F_Primer: AACAGGGTAATGGTACCTACTATATCGCTAATGCTCTA
LHA_R_Primer: GCACTACGCGGCCGCATATCAGTTTTGTTGTCGAAGAGTGCGAA
RHA_F_Primer: CGGCCGCTATAACTAGTAACAGATCTAAAATCTATAATCAAGTAAAAT
RHA_R_Primer: CATTACCCTGTTATCCCTACGACATTCGAGGGCTTGGC

***Gene: CHCHD2, Construct: pWhite-STAR_CHCHD2***
gBlock_CHCHD2_LHA:
AACAGGGTAATGGTACCTACATAATTTTTAGAGCCTTAACATTCGTTTTTAAATCTTAAAA
ACGTCTTAAAACAAAAATAAACTACAAAAAAAGTTCCAAGGTTTATGAAAGACGGAAGT
TTCAGTGAAGATTACGTTTAAGTATACAAAAAGGGTCACAATAACTTTTAAATACATATT
TAAGTAGTTTTCCTGCGTTAACACGTTTTTTTAAACTGTAGATGGGCGCTATACGCACAC
ACACACACGCATAAGCACCGTAAAAAACACACGCGCGTTGTTGTTGTTGAATTGCGAGTT
TTTAAGGGTTTATACTTTCTTGAAGGTTATCCTTTTGGATAATTGGCTCAATATACGTGGA
TGTATATTGTATTAAACATATTCTTGCGCTGCAAGCAGATAAAAAAAATATACAAAAAAA
AAGGGGAAACAAACACGTGTGACGCACACACACAGTTGCACTCGCGCGCACACACACAA
ACAAAAAGCCGCCCGTTTTTAACCTCATTTCGGTGGAAATGGCAATATTCCGCAGTTTGC
CGCAGTAAATTCACAAAACCCTTAGTATCCACACTAGTTGCCAGCGTTTAGCCGTCTTTTG
GCCATCCTCTGGCCACAATCCACACAATTAAACGTTGAAAATCCCTTACTTTGTCACAGA
AGAAATCGAGATACGATGTCAGCGCAAAGGCAAACGCCTGAAAAAGAACATGTTACGGC
AGTGTGCAGCAACGCCAGAGAGTCCGCTTCCGCCGACACGCACACGGTTGCATTCCGGCT
GCCAGGGCTGCGCAACAAGTGAACTATCGAACGGAGATAGGCGGCCATGAGTGGGGATT
ATTCCTCGACTTAATCACTTTTTCGAACGCGTTTGCTACTTGAGTTGCACATTTATATGTAC
TTCATAGAAATATACCCTGCTTTCAGGCATACTATTAGTTTTAAGCCACTTTTTTAATGAC
AAATTTCGAACTTTCTATCGATATGCGGCATTTCGCGGTTTTGCCAGTCACAACAAATGGC
GATAGTTGAAATTCGCATGCCAAATTTTCTTTATTACGGTCACCTTATAGAAATTTTGATA
TGCTTGAGTATTAAATGTTATTATTTCACTGACTGTAAAATAAAAAGTTATTTAAATCGTA
ATGCATATATATTTATAACATATAGTTCTTTATTTGATTTTGTAAGTGGAAGATATGCGGC
CGCGTAGTGC
gBlock_CHCHD2_RHA:
CGGCCGCTATAACTAGTAACGGTTAAACGAGCGATTAGTCATAGATCTTATTAATGGTCT
TATGAAATATTTGCCGTGGAAACGTAATTTTGAAAAGTTAAGAACTGTTTAAAAGGTTCC
CTAAATTACCACCCAAAAAAATATTATGAACTAATCCCTAAATTCGAGACATTCATGTTT
CCGTTTTTATTCAATAATTTTTACATAGAGTCATCACTGTTCATAATTTTGCAGCGTATTGA
TTTTTAGGATTAACAATTTTTTTGTAATGGGGCAATACGTATGTAACAACTACAAATTTTT
CGACTCTAGTAACACTGATGTCTCCTATTATCGAAAAACAGTGTCTATATATATATATATA
TAAAAAAAGAAACAAAGAGTTATAAAGAACCATCAAAATGCCAAATGGATGAAAGCTAT
CGTTTTCGTAGCTTTTTTGAAGATTTTTGGAATTTTCGATTCTGGTCGTTTTCAAATGGTAC
GATTTTAGGGGATTAGAGTTTTCCGAATTCAAAAAAAAGTCTCCGGGGAGGAGCATCGAT
CCACCAGGTGTGTATGTGTGTATATATAATGTTGCATATTTGTAGTGTTCACTGCTCAATT
TGGAATTCCGCTTCAAATGTGGATTTGCCTTCGATTGTCTGGGTTTTATGGGTAAAAACGG
ATGATATAAGGGGTCATCATCTATCTTCAGAGCTTTCCGGTCTTTGTGGTCTGCGGGCCAA
GACTTTCCCGGGACTTTTTCTTGTGCAGCACGCTGGGCACCTGAACCTTTGGACTGACCAC
CGGTTGCTTGGGTGGCTTCACCTTCTTGGATGATCCCGCCGCCGCGGCGGAGCTGCGCTTG
GCCAGGAAGTAACCGTCCGTCTTGTCCTTGGTGATGCACTTCTTGCCGGGATATGCTCTCC
TGCGATAGGGGGTGATCGCGTCGTACATTTTGCTTCTGACCACATACGCCTGCTTCCAGCT
TGAGGCCACCTCCTTCTCTTTGATGCGTGGCTTGTCGAGATCCTTGAAGGGATCGGCCGCC
TTGCTAATGTCCGCTTTGCTGGTCACATGAGGCAGTTTCTGGCTCGCCACTCTGTGGTAGT
CAGTGGCCGATGATGGGGACTTCGGTTCGGATCTGGGCTTGGCTTGGGCCTTGACGATGG
CCTCCGTAGGGATAACAGGGTAATG

***Gene: Coq2, Construct: pWhite-STAR_Coq2***
gBlock_Coq2_LHA:
AACAGGGTAATGGTACCTACTTACAGCGGTATAATTTAACCCATGAATCTTTATTCATGA
GACACCCTGTTCGAGCATATTCAATTTGTATTATTTGTATTTGCTGAGAGCGAATAAAGCG
CTAGAGAGCGAATAATTCGTTGTGGTTTCTGGCACATAAACATACTTATACGAACTTATA
CTAGGGAGTGACTAGCCAGAGAATTCAAATTCCAAGTCGGACGAGATATTTGTTTGCTCT
CCGTGCAAATCTCATTCAGCTAAACTGGATGAGTGACTTTTAAATTACAGCGTTTAGCGC
AGTGGTAAGTGATTTTGTATCAAAATTGTTGGACACAGGAGTTGTGTAACTGCGTACTCA
CCACCATGATCATAGGGCAGGACTTGGCCGTGCTGGTGCTGTCATCCGAGCCGTGATCCA
TCTTGGGCTGCTGGTGTAGTATAGATATAGATAGAAGGCGTGGAAAATCCGACTGGCGAT
CGGGGAGGCACAGTGATAAGAGCTAGACCTGCTGAGGTGCTGAGAGCAGAGCGTTGACT
GCTTGAAACCCGCTGCTCGCGAGTGGTTTTTAAAGAAAATCGCACCTAATCCCCCGACG
CGGCGATAGGAAAGTGCGCTTATCTATGGGCAATGGGAAAGAGCGTTATCACCCAGTAG
CGGCGATACCAACGATAGTCGCCCCACACTAAGAAAGGCAGCCCAGATACAGTCATGTA
AATAAACATAAATGCATGCAGATCTCAGTGTTAGAGGTCAGATCAAAAGATATATTCTGG
GTATCAGTTGCAGATCTTTATCTCGAGAGATAAAATCCATAATTGATGCGTCGTGGGTTTC
TCTGCaaTTTGCAGCTTAATTTATGGACTCATGGCAAAGGTTCGACAATCGATTCGTGTGC
AGAATTTATTGCAATTCAGATTATCAATCATGGGCTAATACAACCCCCTGAAATTGTTGCT
GTAATACATGATGTTGAAATAATTCCATTATTTCTTCGCTTCGATCACGGTTTTATGGCTC
TAGTCGAGTACATTCATACGTAGGTATATGCCCATATTCCGGCATGGGAGCGATAGCAAC
CGTCGGCCGCCGGATCGCCGGAGGTCTGCCTTAGTCCACTTGGCTTTCCTAGAGTGTTATT
TCTTACTGTGCTATTTCTAACTGGGATAGAATGCCGCCAAAATTGATGGCCGTGCGCAAA
CCTTCTGCGGCCGTGTGCATAAATTCTACGGACATTCCTGCGCCGTGTGCCAATTTCGAAA
AATGGGCGACGTGCGCAAAACATTAACGATATTGCAGAGGGGTAAATGGTTGGGTCTAC
GCAAAGATATGCGGCCGCGTAGTGC
gBlock_Coq2_RHA:
CGGCCGCTATAACTAGTAACAATTGTTATATTACCTTACAGAAAGTCACTTATTCATCCAT
TTGGCATTTCAGATCTATTCCCTCAACATTGACAACCCCAGCGACTGCGCCAAAAAGTTC
ATATCGAACCACCAAGTAGGACTCATTCTCTTCCTCGGCATTGTTCTGGGCACCCTTCTGA
AATCAGACGAAAGCAAGAAACAGCGACAATCCTCACTAACAACATCGACGGCCAGCTCG
TACGTTCCAGCGCTGCCGCAAAAGCCAGAAGTTTTAAGCTGAGATGAGCCAGACGAGCG
TAGATTGTAGTGTTAAATTATGTTTATAGCGCGCTTTTTGATTCGCCAAAACCAAAGTCTC
TAGGTATCTGATTAGATAAAATCTTGTAAATAGGCTTATGACCATTTTTAGCTCATTTGCA
ATAGGTTAAGTCTTAAAGAATATTACTAAGTGCACACCTCACTTAAACATCAATTAATTCT
TAATTGTAAGTGCCCGCCTTGACGTAAGAACTTACGTAAATCACCCATCCCCTGAGCCCT
CCTAACCCATTTGAAAAGCAAACTGGTTTCTGTATTAAATTCTTACGTGTTATTTATTATT
ATTATACAGATTTTGGTTGTCTATTGAAATTCAAATACTCTTCAAATGTATTTACTTCATTC
GTTTGTTGTTACGGCTAGGAAAAGTGCGTTACATAAAAGTAATGATAGAATATTAGTTTA
GATCAAGTCAGCCACGTTCATGGGCATCTCGTCGATTTGTGTGGAGTAGTACTGCTCAAT
GTCGCGCAGGATGCGGATATCGTCCGATTTGACAAAGTTAATTGCAACACCTTTGCGTCC
GAAACGACCAGAACGACCGATGCGATGGATGTACAGCTCACGGTTGTTGGGCAAATCGT
AGTTGATGACCAGCGACACCTGCTGTACATCGATACCCCGAGCCCACACATCGGTGGTGA
TGAGCACTCGCGACTGGCCGGCTCGGAACTCCTTCATGATCTCGTCACGCTCCTTGTAGG
GATAACAGGGTAATG

***Gene: Dj1-a, Construct: pWhite-STAR_Dj1-a***
LHA_F_Primer: AACAGGGTAATGGTACCTACCGATTAAATTACATATATGC
LHA_R_Primer: GCACTACGCGGCCGCATATCGGATGGGTACATTTCGTGGG
gBlock_Dj1-a_RHA:
CGGCCGCTATAACTAGTAACCCCCTTGTTATATCATTTGCAATTTTAGATCCTTGTTACCG
TGGCTGGTTTGCATGATTGTGAACCGGTGAAGTGCTCCCGATCTGTGGTCATCGTGCCGG
ATACTTCACTGGAAGAGGCCGTGACCAGAGGTGACTACGATGTGGTTGTTCTTCCTGGCG
GATTAGCTGGCAACAAGGCGTTGATGAACTCGTCTGCCGTTGGCGATGTGCTGCGTTGCC
AGGAATCAAAGGGCGGCTTGATTGCCGCCATTTGTGCCGCTCCCACGGCTCTCGCCAAGC
ACGGAATCGGCAAGGGGAAATCCATCACTTCGCACCCGGATATGAAGCCCCAGCTGAAG

GAACTTTATTGGTAGGTTTGCCAGTATTACGTGTTTATTAAATTATCAGGAAACTCTGACT
ACTCAGTTATATAGACGACAAGACTGTGGTCCAGGATGGCAACATAATTACAAGTCGTGG
TCCTGGCACTACTTTTGACTTCGCCTTGAAGATTACCGAGCAACTGGTCGGAGCTGAAGTT
GCCAAGGAGGTGGCCAAGGCAATGCTCTGGACTTATAAACCATGATGGGAATCGAAGGA
AAAGCTGATCATAAGTTACATAAAAATAAAAATATGAAACATTAACGTATAACGTATAGT
TATTTGCAAATAGTAAAATCTCTACTATTTTACAACCAATGATGCTGAAAAAATCGGTGT
ACTGATTTTTATGTACAAAGTTCCATAACTAGAATTTTCCTATATAGAATGACCTAGGTTA
TTGTCAATTAAAATATGAATATGTGGAAATTCATTTCGTATCTTTTTGATTTCCCTATTCAG
TAAACAACTGTACCTGTACCTATTTCCGCCATAATATTATCGAAGTTTCCCACCCAAATTT
TATATCAGAGTTTCCCATTTTGCCCGTTGGCGCTTTATGCACACCTTAGGTAAGACCGTAA
TACCGCCTAGTGAGACCATGCGAGGACTTAAGAATAACACCACCATACCGGGCGATTGC
AAATGTATTGAAATAAAACTAAAAGAAATCGTATCAACTGCTAGTTCGGCGAAACTCGG
GAAATCGGAAGGTTCAACTTTTGGGGCGGTCCGTGAATTTTTGCATTTTTCTGTGTTTGG
CGGAAAACGAAACCGTTATTGAGATATTTGTGAATCGCAAACTGTAGATACAACTTTATA
GAGTAGTTACAGTCGGTAGGGATAACAGGGTAATG

***Gene: Dj1-b, Construct: pWhite-STAR_Dj1-b***
gBlock_DJ1-beta_LHA:
AACAGGGTAATGGTACCTACGATTGCTTTTCGTGCGTTTTCCGTGCAGTTTTCCTTGCTTTT
CCACACAGGCAATTGGAAATGCACACCAACACACACATGGGGCGCACACCAGCGCAGCC
ACACACGCACTGGCTTACACACACACACACTCGCACACATTCAACAGATATTGCATTTTC
TTTTTGGACTGGAAGATACGCAAATTCCAGCTGCAAACGTTGCATTTTCACCAGGCATTTT
CCGCAATTACCTTCTTCGCGTTTTCCACGTTTTCCACGCAATTTCCTCGCTTTTCGCCTCTT
TGTTCTCGCCTTTTTCGCGTTTCCCACACACGGCACTTATTGCTTTTCCTCTTTTTCTCCGTT
TTTCCTTGCCTTTTGACGACAATTCGCTGATATTTGCCGACGTATTTTCACTTGATTCACTC
GCGTTCGCAACCCATCACTTTTCGCGTGCATTCAATCGCATTTTTCAGGCATTTCCGATGC
GCGTTTGGCTTAAATTTAGTAAAATTAATTCGTGGTTTTGTTGCTGCCGTCGTGCTAGATA
TGTTCACATATCGATAGACCGATATCGATAGACAGAGATCTAGGGAGGAGCAGCTGTACT
TGGTCTGGCAACCACATCAGGTATCGATAGGATTTGACCGTGGTATATAAACATCGATAT
TATACTGAAAACAAATATAAAAATATTTCAGTGATCAGTTTATATAAATTTATGTAATAC
GACAGAATATAGAGAGCAAGTAAATGTTATCGGTTTTAACTTAAAGAACAAGACTTGTTA
GAATTGTTAATGAGGTTTCAAACCATTTCTGTGTAGCACGTAGCATAGACGCAATCCTTTT
CTCATATAAATAAAATCAACTGATTTCAAAGTCTTAATATTTTAAAATATAATGCAGAAC
ATTGTTTTGAGTAAGAATGAAAATATTAACTTCCTGTGAAAAACAGAAGTTTTAAATGCA
TCAAATTAGTCGCCTAACTGCAAGTCAAACCAAATTATACAAGTGTTTTAGTTTGCCGCTC
AGTTTTTGAAACCTCGGATATGCGGCCGCGTAGTGC
gBlock_DJ1-beta_RHA:
CGGCCGCTATAACTAGTAACATGATAGATCAAGGTCACCGTAGCCGGTTTGAATGGCGGG
GAAGCGGTGAAGTGCTCGCGGGATGTGCAGATCCTGCCGGACACCTCGCTGGCTCAAGTT
GCCTCGGATAAGTTCGATGTGGTGGTGCTGCCCGGCGGACTGGGTGGCTCCAATGCCATG
GGGGAGTCCTCGCTGGTCGGTGACTTGCTGCGCAGCCAGGAGTCTGGTGGCGGACTCATC
GCCGCCATCTGTGCCGCGCCCACCGTTTTGGCCAAGCACGGCGTCGCCTCCGGGAAATCC
CTCACCTCGTATCCCTCAATGAAGCCCCAGCTGGTGAATAACTATAGGTATACCACCTCTT
AAGTTTATTTTCCCATGATCTTCATCCCATTTCTTTTGATTATTTCTTTAAGCTATGTGGAC
GACAAGACGGTGGTCAAGGATGGCAATCTGATCACCAGTCGAGGTCCTGGCACCGCCTA
CGAGTTCGCCCTCAAAATCGCCGAGGAGCTGGCGGGCAAGGAGAAAGTCCAGGAGGTGG
CCAAGGGTCTTCTTGTGGCCTACAACTAACATATTTACACTTATTTATGAAAACAGATTGT
TTCAAGAGCAACTCATGCAAACAACAACTGAATTAAAATAAAATTATTCAAGACAGAAG
TTGTGATTCAATTTTATATTTGCTATTTGAAAGAAGTTTAAATGGAGAGCTAGGAAATGG
CATGTTGTTAAAGATATATTAAAGAAATTTATTTAAGTGAATTTAAATGTCATTATTTGAA
GCAAACGGTTTTCATTTAATTAGTTTTGAGCGAAGTATAATTCTATCATAATGAATCAACT
GAACTGCGTTAAAATAAATTGTTCAATGGGCTTTAATTTAATTTGTCATGAAATATAAATG
CATATATGGCATGAAAAAAGAAACATCTATATTATTTCATTTGGTTATGACTGGACTATA
AATGAAATTAAAAATGCATATATTAAAATGGGATGTGAAGCCAAATGGAAGTATTGCTG

ACGATATATTTACAAAATAGTACATTATTAATATATTTAGTTATATTCAATATTTACATTG
ATTAAACATTCTGCTGCGTAGGGATAACAGGGTAATG

***Gene: rme-8, Construct: pWhite-STAR_rme-8***
LHA_F_Primer: AACAGGGTAATGGTACCTACAATGGTAGTTAAGTTCAG
LHA_R_Primer: GCACTACGCGGCCGCATATCCAATTCATGACTGGTCTT
RHA_F_Primer: CGGCCGCTATAACTAGTAACTTATAAGATAATACGGAGAGCTCGGAACT
RHA_R_Primer: CATTACCCTGTTATCCCTACAGGCAGAGTATACGCTTC

***Gene: auxilin, Construct: pWhite-STAR_auxilin***
gBlock_Auxilin_LHA:
AACAGGGTAATGGTACCTACCGGTGGTAATTAGTACACTGATAAGGTTGTGTATTTTGCA
TTCGACAAATCTCCACGCGTTACGTGAAACTGGCCAGGTGCACACAGCAAATTGCAAGCT
TAACTTCAATGTCCAAAATAAGCGGAGCTCGGTTTGCGCAAACTTACGGCCATATCGGTC
ATCTGTAACTATCCGTATTCGCCAGCGGCCCAAGCGTAGCGCAACACAGCTCTCAATTAT
CCAAAGACAGATGTGTAAATATACTGCCGTTACCAACTTGTTGCTGTTCCACGGAGGGGA
GGTGCCCCAAAGGCTTACTAGTCATACTCCATTTCTTCATTCCGACGGCCAGCTGTTGCAG
CGGTTGTCCCCTGGGTTTCCCCTTAGCTGTTGACTTCAATCAGCCAACTCTACCTGGCACA
ATGTACATAGGAAACTCAAACTTCAGAGCTGGACCTTGAATTGGCTCAAAGACACATGCA
CGCTTGATGCACAGCTAAATGCGGGAATTTACTCAGGCGACTCAAAGGGGAATGACTTTT
CGAAAAAAGTGTTTAACCTTTCAATTACATTTTATTTTCTTTCAAAGCTGTCACAGCCCCA
GTCAAAAATGCAATACCTCTAGACCTCTGATAAATGTGGTTGTGTGCGCTCTTTTAACCGT
ATCAAGCTAAATGGGTACTTGTTGAAGAGTATGAACTAGCGGAAGGAACTCTTTAGAGCA
TATATCTTGACATGACTGCTCGTACAAAAACTGTGATCTCTTAAAACCTCGCAAGGGCGT
TGGCACTGCACGCATTTTAAATATAATACAATCACCTCAGGTGCAATCTTAAAGCCAGGA
TTAAACGTCTGAATTTTGAAGAGTCAACATACAAAATGTTTGCTTAAAAGCACTCTAAAT
TTAGTCGAAATTCATAATCTAGTTATCCGCGCGCCACATCGATAAGAGCCCAGCGTCAAA
TATCGATAAGAGCAGATATGCGGCCGCGTAGTGC
gBlock_Auxilin_RHA:
CGGCCGCTATAACTAGTAACTCGAGAGGCGCAGCTCTGCTTGAGCCAAGCCAACGGACG
AATTGTGGCGATAGTGCATTGCCACCATCACGGTTACGCGTTACCGCTGGTGCCCACCTG
CGAGCCATGGCACATTAAACCGATCGGGTTTTTTTTAACAAGTTCTTATATAAGAACAA
CAGTGGCCTGAAGGTGGCTGAAAGTTCGTAGCTCTGCAGGGAGCTGGAAGACAGAATTC
GTAAATATATAGCAGGCTACCAGCCGGTCAACCCCTTCCTCCGAGCGTGCCGCACTTAGC
ACTTAGGCAGGGCCTTATCTGTGCCTCGTTATCTACTTGTCTCTCGCTTCCACCCCCTTTTT
CAGGCGGATACGCCTTCGTGTATGTAGCCCAAGATGTGCAAACTGGCACAGAATACGCCC
TCAAGCGTCTGATCGGGGCTGACATGCAAGCCTCCACCGCCATCATCAACGAGATCAACA
TCCACAAGCAGCTGTCGGGGCACGAGAACATTGTCGCCTTTGTGGGTTCCAGTTATACCG
CCCCATCAACTCAATTGGGAGCCCAGTACTTGCTCCTCACCGAGCTGTGTAAAGGTAAGG
GACGCCCTAGGGCGTCCAATCATACTACGTGAACATTTGAGCACATGGATTTCTATCCTT
GTCAGGCGGATCTCTGGTGGACTGCTTCAGAACAAACAATGCTCCATTCAATCCGACTTG
TGTTCTGCGCATCTTCTACCAAATGGCGCGGGCTGTGGCCAGCTTGCACTCACAGTCACC
GCCGATAGCCCACCGAGATATAAAGGTTGGTCTCCAACCATAGCTGGAAGTAACAGACT
AATACCCTAATCCTATTTGATTGCAGATTGAGAACTTTCTCATTGGCAACGACAAACAGA
TCAAGCTGTGCGACTTTGGGTCCGCCAGCACGGAGGTCCTGTCGCCCACGTTTGAGTGGA
GCGCCAACCAGCGCGTAGGGATAACAGGGTAATG

***Gene: eIF4G1, Construct: pWhite-STAR_eIF4G1***
LHA_F_Primer: AACAGGGTAATGGTACCTACATGGCCGCTACGCATCTT
LHA_R_Primer: GCACTACGCGGCCGCATATCGTTATTTCACATTTTTAAACGCTT
RHA_F_Primer: CGGCCGCTATAACTAGTAACCATTCAGCACAAAATATG

RHA_R_Primer: CATTACCCTGTTATCCCTACATATTTATTTTACTCGGC

***Gene: ntc, Construct: pWhite-STAR2_ntc***
LHA_F_Primer: GTCTCTAATTGAATTAGATCCCTGTAAATATGTTTAAATCT
LHA_R_Primer: CGGCCGCTATAACTAGTAACACTATTTTAAACAAGTTGGAACTGA
RHA_F_Primer: GGCACTACGCGGCCGCATATCTAAAAAGAATATTCCTTGCTTTATT
RHA_R_Primer: TAACAGGGTAATGGTACCTACCGAGCAGCAAGCATCCCGC

***Gene: Gba1a_1b, Construct: pWhite-STAR2_ Gba1a_1b***
LHA_F_Primer: TCTCTAATTGAATTAGATCCCAATGAAATGATGTTCAACA
LHA_R_Primer: GCGGCCGCTATAACTAGTAACCTATTATCTTACTTCTGTCCAGTCG
RHA_F_Primer:
GGCACTACGCGGCCGCATATCAGAAAACAATAAATTCATGCCTAAATGGCCATGAGACTC
CTCAA
RHA_R_Primer: TAACAGGGTAATGGTACCTACTGCAGATCCCGCGGATCGA

***Gene: punch, Construct: pWhite-STAR_punch***
LHA_F_Primer: AACAGGGTAATGGTACCTACGTTTGAGGGCAAAAGCTT
LHA_R_Primer: GCACTACGCGGCCGCATATCTCATTGACTAACTAAATC
RHA_F_Primer: CGGCCGCTATAACTAGTAACACTGATCAAGACGCCGGA
RHA_R_Primer: CATTACCCTGTTATCCCTACTTGGGGGGAGGTTGACTT

***Gene: Gdh, Construct: pWhite-STAR2_Gdh***
LHA_F_Primer: TCTCTAATTGAATTAGATCCCGGGGTCCATTCAAAACCGA
LHA_R_Primer:
CGGCCGCTATAACTAGTAACAGACGAAATGCCAGAGTAACAGCAACAACCGGCTTGATC
AGTC
RHA_F_Primer:
GGCACTACGCGGCCGCATATCTAATAATAATATCTACATTCTTGGGCACCACTCCCGGCT
GGG
RHA_R_Primer: TAACAGGGTAATGGTACCTACATGATACTTCAAAATTATA

***Gene: Omi, Construct: pWhite-STAR_Omi***
LHA_F_Primer: AACAGGGTAATGGTACCTACAGAGGAGCGTCGAGCTAA
LHA_R_Primer:
GCACTACGCGGCCGCATATCAGACCACATTCGGAAGTATGTTTTAGCTATTCAGCTATA
CGTTttAACTAGAAACGACAGCA
RHA_F_Primer: CGGCCGCTATAACTAGTAACACGTAAATCAGACATCCGATCTGG
RHA_R_Primer: CATTACCCTGTTATCCCTACCGCACCCGATTAGGAATA

***Gene: Lrrk, Construct: pWhite-STAR_Lrrk***
gBlock_Lrrk_LHA:
AACAGGGTAATGGTACCTACTTGGATATCTCGTGAAGTACACAAACCAAAACAATGAGC
GACTCCGACGAAGGTGAGATAACACAACTTTTAGTGAAATTATGCACAATATGCTAGTTT
TGCGCAAATTTAGTCAGAGTGCAAATGAGCATTATACGCTTGACTATGCAAATTACCTAA
GTGGTGCAATTAAAACTTCAACCAGTTGTTATTGGTTTTTCAACAACACCAGCAAGGGCA
CAGGGTGACATTGGTGCAATGGAAGTGGGCGCGCAAAAACTTTTTTCAAACAGTCATCCC
CCACAGAAGCAACTGGTGCAACCACTGGACCATCCGACACCCACGAAGAGACAAATCAA
AACAAATGCCACGCACTCACCCAATGCCACATTCTTGAACAAGTGCAACAGTTGCCGCGC
TTCTTGGGCTCCGACTTGCGCAGTGCGACATAAAACGGCCTGGATCAATGACCACAAAAA
CGTGGCCATCGATTTTCCGCAGTTCAGTTTTTGCTAGCGGCGCGTTCGGTCATCATACCGA
TGCCAATTAATTTTTGGGGCTTATTCGCACCGATTTGGGTTACATAATGTCATGGAAATTC
AGCATGGAACATCCCAAAACTGGAACTGAAACAGCTCTGGAAGCCTGTGACTACTTCGTG
GATGAGGTCATTGAGGCATCCTCAATACGGGGTGGGAATTGGTTTATTCCAAAGGTGCTG
TGGATATGCGGCCGCGTAGTGC

gBlock_Lrrk_RHA:
CGGCCGCTATAACTAGTAACGTTTTAGAGCTTGTCAGTCTGGTTACGAGAGCATTACACA
GCGATTGCTGGATGCTGGAGCGGATGGTCGCTCCCATGCCGTGACCAAGTACTCACCCTT
GTACGCCGCCGTCCATAGCGGTCACTTGGGCATTGCCCGACTGATGCTAGACCATTTTCC
AGAACTGATCCAGCAGCCGACTGTAGAGCGCTGGCTGCCGCTGCACGCCGCCTGCATCAA
TGGACACATCAAGCTGCTGGAACTCCTTATCAGCTACAGTTATCCCGACTACCTCTACCA
GACATATCGCGACGAGGAGGGCCAGTGGAATGGCGGCTCCCCTTCGACGCTAACGCTC
ATGATGTGACGGGTCAGACGAGTCTGTATATTGCCAGCATCCTAGGAAACAAGCAGCTGG
TTGGTGTGCTTCTAAAGTGGCAGCTCCATTGCCGGCGTACGTTGGGCGATTCCGCCAGCTC
GGTGAGCACTCCTATTACGCCCACCAGGAAACGTATTTCATTTGGCATTCAGGCTATCAT
GTCCAAACTGCACATATCTGGAGAGTCAGAAGGACCCGACGACCTAGCTTCACAAGAGT
CAACCGAGTGCCAACGGTGTCCCATTAACGTGAATCTGCTATGTGGAGCGGCGAGAGAA
ACAGCTCTGCTTGCGGCCGTTCGAGGCGGCCACCTAGACGTGGTGCAGTCTCTGCTACAG
CACGGCGCAAATCCGAATATTGTAGCCAAGCCAGTTGAGGATCACAACGACCCGAAATG
TTGCGAGGAAATATATGGGCTCAGTAATGTCCCCATTGCAGAGGCCTGCAAGCAAAGGTC
GCTGGCTATGTTGGATCTCTTGCTTAAGCATGGAGCCCGCGACGATAATGGCACGGCCAT
TGGAATGGCTATTACATGCGGCGACGAGGCCATCCTGAGTCGTCTTTTGGCCCGACGAGT
CCATCCGGACTCAGACTACAAGATCAACAAGAAGGGTCTTCCCACACCAGTGGAGGTGA
ACGTGTTTTTGCCGTCCACCAGCAACGTAGGGATAACAGGGTAATG

***Gene: Tango14, Construct: pWhite-STAR2_Tango14***
gBlock_Tango14_LHA:
TCTCTAATTGAATTAGATCCCGAATCCGGCGCATGGTCACACTGACGAACACGTATTTTC
ATTATTCTTGCGAAGCGCATCGCTTAAAACTTGATTACGCTCGGATAACAAGATGCACCC
GCGTCTCAAGTGACTGAACAGGTAGCGAAGTGACCCTGGATGAGCGTGGGTAGGCAGCC
GGCACCTCTGGAAAGCCCTAATCCCAGGTCCGCCAGCCGTTTGCAGGTTATGTCCGCTTG
TGTACACTGTGACGGGGAACCGAAGAAGCAACAGGAGCGGATTCCTGGCAGAAGAGACA
ACCGCTGTACGAGATACCGCCCCTGACCTGCGCAACATGATCGAGGTGCTGTGCCTGCTG
CTGGGCAGGATCCTGCTGCTCCTGGTCGGCGGCTACGAACTGATGTGGCGGCTCCGAGAG
CGACTGAATGCACTGGCCGTCCGGTCATATGACCTTTGGCGCAGCAAAGCGGCACGTGAG
GCGCACGAGCGGCGCGTGCTCGCTGACTGCCGCTCCCAGTTGACGAAGACGCCGCAGCAT
CTGGTCCTGGTTATCTCCCCCGTAGATGCCGGCGTGGATGCGGTGCTCCTCAGCAGGATCT
TCGACTTTGCCCTGGACGTGGGCATCAAACACGTCAGCCTGTACGACAGGCGGACGAAA
GGCAGGGGATACGTGGACATGGCCGATCTCTGTCGATCCACCAACGCGGACACGGGCAG
CTGCTTAAAGTGGCCACCCGTAGCCAGTCCCAGCAAACTGGAGAACCAGCCCAAAAACG
GACAAAAGACAAATGGTTATGTGAACGGTTCACATTCCCCTCAACTGCAGGTGAGTGGCG
GCGTCTGATAAGCCGTGATTAGTAATGGACTTTAGAACGATGTCGGAAAATGTCTTACAA
GTATTACACTTTTTGTTTCAATTGTTAAATAAAAGAAGTAATGAGCTGGAGATTTAATTTA
GTTCATCGTTACTAGTTATAGCGGCCGC

RHA_F_Primer: GGCACTACGCGGCCGCATATCGTAGCGACACCCTTATATT
RHA_R_Primer: TAACAGGGTAATGGTACCTACCCTCGACTAACGAGAGCAA

***Gene: Park, Construct: pWhite-STAR_Park***
LHA_F_Primer: AACAGGGTAATGGTACCTACCAGACCAAAACTGCATTAC
LHA_R_Primer: GCACTACGCGGCCGCATATCGGCATGTATTTACCATATGC

gBlock_Park_RHA:
CGGCCGCTATAACTAGTAACTGTCTTCTAGTAGCAATGTGACTTGGGTCAGCAAAGCGTT
TTGCATGCCATTCGTTTGCGACCGCCTGTCCAGCGCCAGAAAATCCAGTCGGCCACTTTG
GAGGAGGAGGAACCTTCGCTTAGCGATGAAGCCTCCAAGCCTCTAAATGAAACTCTGTTG
GACTTGCAGCTGGAAAGCGAGGAGCGTCTGAATATAACCGATGAAGGTAGGCAACAATT
AGTTTTTATTGTTTTTTTTTTTAACGAAACCCCTTGTAATCCCGTCGTATCCTCTCCAGAAA
GAGTCCGTGCCAAAGCGCATTTCTTTGTACATTGCAGCCAATGCGATAAGCTGTGTAATG
GCAAACTTCGGGTCCGCTGCGCTTTATGCAAGGGCGGCGCCTTTACTGTCCATCGTGATCC

GGAGTGTTGGGATGATGTCTTGAAGTCGCGTAGAATTCCCGGTCATTGTGAAAGTCTGGA
GGTGGCCTGCGTGGACAACGCGGCTGGCGATCCGCCCTTCGCTGAATTCTTCTTCAAGTG
TGCTGAGCATGTCTCCGGCGGGGAGAAGGATTTTGCGGCTCCATTGAATCTAATCAAGAA
TAACATCAAGAATGTTCCTTGCCTGGCCTGCACGGATGTGAGGTGATTATGCAATAAGGA
ACATCTTAATTTCTTATAATTTTCATATCATCGAAAATGTCATAAAAAGTTCTTTTAAAGT
CTTTAAAGTAATTATTTGTCATATAATTTTATTCTGCTATAATCACCTTATTATGTTCGCAG
TGATACCGTGTTGGTTTTCCCCTGCGCATCGCAGCACGTCACCTGTATCGACTGCTTCCGC
CATTATTGCCGTTCCCGTTTGGGCGAGCGTCAGTTTATGCCGCATCCGGACTTCGGCTACA
CCTTGCCCTGTCCCGCAGGCTGCGAGCACTCGTTCATCGAGGAGATTCATCACTTCAAGCT
GTTGACACGCGAGGAGTACGATCGGTACCAGAGATTCGCCACCGAGGAGTATGTCCTAC
AGGCAGGTGGAGTACTCTGCCCCCAGCCAGGATGCGGCATGGGCCTTTTGGTGGAGCCCG
ATTGTCGCAAGGTGACATGCCAGAACGGTAGGGATAACAGGGTAATG

***Gene: Pink1, Construct: pWhite-STAR_Pink1***
gBlock_Pink1_LHA:
AACAGGGTAATGGTACCTACGGCGGCTCCAACGGGCACTTCGAGCTAAATGTATTCAAAC
CCCTGGTCGTGTCCAATGTGCTGCGCTCCATTCGCTTGTTGGGTGAGTCGGAAATGTGTTT
TCCAAGTGGAAAAACATGCATACATAACCTTCTCCTCCCAGCTGATGGCAGCATGACCTT
CAGCAAGAACTGCGTGGAGGGACTGCAGGCCAACAAGGAGAGGATCGACAAGATCATG
AACGAGTCCTTGATGCTGGTGACCGCGCTGAATCCGCACATTGGCTATGACAAGGCCGCC
CTGATCGCCAAGACGGCGCACAAGAATAAGACGACCTTGAAGGAGGAGGCACTGAAGAC
CGGAATTACCGAAGAGCAGTTCAAGGAGTGGGTCAATCCCAAGGAGATGCTGGGACCGA
AGTGATTCGCTGTGCGGCTATTGACGATTTTGTTTTAGTTTCGTGTAATAAATTTTTGTGTG
TAGCAGGATATTTCAATAGATTGTATTAATTTGTTGATTTTACGGCAGGTGAATAACGGTT
TTTAGTTTTTGAATTCTCCAAATGATTCGAGTCCACACTACGCGATAGTCATTATCGATTC
TAAAGATATACCGTAAGTGTGGTCACACTAACGCCTTGCTGTTTCTAAATTATTCCGTTGT
TTTTATCGTAAATTCTTGCACACTTAGCTATTATTTTATAGCGAAATATAAGCAAAGCAAG
CTTTTAAACAAGTGTTTATTGCGGCGCACATTTTCCGTGACAAATGAAAAGATCCAGCTG
ACCAGCGATTATCTAACGACCAAATCGAATCGAAACGAAACGTTGAAGTAGGCGCATTA
CATTAAAAATTGGCTGCCAGGCAAGTTGGAGGTCGTAAATTAATCGTTTTGTTCAGTTGA
AAATAGCCCAAAATCAGTCTAGCCCACACAGATATCGAAAAAGGCGATATATCTTTGGAT
AATCATTGCAAAAATAAACCCTTTCGATTCGACGATTTCGCCTTGTAGTTCTGCCACCACC
GCCCCCtCAaTTCGCCCCGCAGCTGCACCACAACCACCACCACCCACAAACAGAAAATTTA
TTTGCGACGCAAAAACAAAACAATCGAATCGAAACGAACCGAACCGAAACGAAACGACA
GCAGCTGTTTGGTTATAAATCGGCGCTATCGATCGCACACATGAGCAAAATTGGCGGAGA
AGAAGAGGAAGCAGCAGCAAGAGTTAAGCACCACAAATCTTAAAGAATAGGTAAGATAT
GCGGCCGCGTAGTGC
gBlock_Pink1_LHA:
CGGCCGCTATAACTAGTAACCATCCAATCTCTTCGCCAGACAACGCCCCAGCAGGCGGCC
AAATCGGTGGTCAATGTAGTGCCCCGCACCATCAACTCTCCGTCGGGATCACCGTTTAAT
GGCAGCGGCAGTAGCCCCACCAGCAGCAGTGGAATCTTCCGAGTGGGCCAGCATGCGCG
CAAATTGTTCATCGACAACATCCTCAGCCGGGTGACCACCACCTACTCGGAGGATCTGCG
CCAGCGCGCTACCCGCAAGCTATTCTTTGGCGATTCAGCGCCCTTCTTCGCCCTGATTGGC
GTTAGTCTGGCCTCCGGATCGGGTGTGCTCAGCAAGGAGGATGAACTGGAGGGCGTGTGC
TGGGAGATTCGGGAGGCAGCTAGCCGGCTGCAGAACGCCTGGAATCACGACGAGATCTC
CGATACGCTAGACAGCAAGTTCACCATCGATGACTTGGAAATCGGTCCGCCCATAGCCAA
AGGTTGTGCCGCTGTCGTCTATGCAGCGGATTTCAAGAAAGATGTTGCCTCGGATGGTGC
ATCCCTGCATACCGATGCGCAGCCGCAGGCAACGCCAGCCTTTGCGCCGAATAGCTGGAG
TACCCACGAGATGATGTCGCCGCTGCAGAACATGTCGCGCTTTGTTCACAACTTTGGCGG
CTCTGTGGACAACGTCTTCCACTACAGCCAGCCATCTGCGGCCAGTGATTTCGTGGGCGC
CCAGTCGAGGGAACAGGACCAGCGGCACCACGAGCAACAGCAGCATCAGAATCAGGAA
CAAGAGCAGCATCAGAACCAGGAGCCCAGCAGCAGTGCCTTCAATGTGGTAAATATGTA
AGCTTAAATCTGCATGGCAACTTGCCATCACGCTCTTTGCTTTAACTTGCAGACTTCTCCA
GCGAATTCAAACATCAACAGCTCTGTGGACAGTTATCCACTGGCACTCAAAATGATGTTC
AACTACGACATCCAGAGCAACGCTCTGTCCATACTGCGTGCCATGTACAAGGAGACGGTA

CCGGCACGCCAGCGCGGCATGAATGAGGCCGCCGATGAGTGGGAACGTTTGCTGCAGAA
TCAAACTGTTCACCTGCCGCGGCGTAGGGATAACAGGGTAATG

***Gene: iPLA2-VIA, Construct: pWhite-STAR2_iPLA2-VIA***
LHA_F_Primer: GTCTCTAATTGAATTAGATCCCGGCTCGAGGAGCTGGACTC
LHA_R_Primer: CGGCCGCTATAACTAGTAACCCAATGGTAGGGGCTTGCA
RHA_F_Primer:
GGCACTACGCGGCCGCATATCTACTACAATGTAAATGGGCTGCAGAAAGCTTGCGATGCC
TTGGCTGATAATCCCTCCTAAACGCTGTCCCATTTGATTG
RHA_R_Primer: TAACAGGGTAATGGTACCTACTCCGGTGCCTCTGGCGGAA

***Gene: Loqs, Construct: pWhite-STAR_Loqs***
LHA_F_Primer: AACAGGGTAATGGTACCTACGTGGCGTCTAGCAAGAACA
LHA_R_Primer: GCACTACGCGGCCGCATATCATGATCGTAAGTTTGTTTA
RHA_F_Primer: CGGCCGCTATAACTAGTAACGACTGGGCTCACGGTCGCC
RHA_R_Primer: CATTACCCTGTTATCCCTACATTTGAAATCACACACGCC

***Gene: Rab39, Construct: pWhite-STAR_Rab39***
LHA_F_Primer: CGGCCGCTATAACTAGTAACAGACAACAAATTCGCCGAGGTGCG
LHA_R_Primer: CATTACCCTGTTATCCCTACTGAATCAACAACATGATG
RHA_F_Primer: AACAGGGTAATGGTACCTACAATCCAAAATAATTAATAA
RHA_R_Primer: GCACTACGCGGCCGCATATCAAAGTAATCGATAGTAGC

***Gene: Synj, Construct: pWhite-STAR_Synj***
gBlock_Synj_LHA:
AACAGGGTAATGGTACCTACCAGAATTTTGCAATAAAATCGCAAACAAAGGTATTAGGA
GTATTAAGAGGCGTAAAAAATTCATTTGAGGAGCCAATAGTTTCGTGAAATCTAGTGTTT
CCTCTTTTAAGTGGTCGCCAAGCATGCCGTTCGTTGGCTGCCTCTAAAAACAGTCAAATGC
CTGTGCAAATTTCTCGTGAAAAATCCATAGAATGCACAAAAAATGTTTATCAGTCACTCA
CAAACACACGCACGCGCCAGCAGTGAGTGGGAGAGAGCGAGAGAGAGCACAGCTGCCG
CCGGTGCATGTCATTGCTGTGTGTGCGTGTGTGGGTGAACTTGAGTGGGAGGCGGAAGAT
ATTGGAAAAATTCCTCCGCTTTTCGGTTTGTTTATGTTTAAGAGTTGCCTATTGATTTTTAA
CAGGCGCCCGAAATTTTAGTGCCATTTCGGTTCAAAGTGTCTGTTGTACTAGTTGAACTTT
CCCATTTCCACTCACCTCGTGCAAATCCTAAAATTAAAAGAATCAAAAACTTTGTGGTATT
AAATTAACCCCAAATTTGAGCTCATTTGAGCTATTACGAAGTGCACAAAATAACTTAACT
TAGCAAGTAAAACACTCACCAAAATATTTTTTAAAACTTAAAAAAACAACGAATGTAGCG
TAGTTGATAATAACTTATTACTTAGTAAAAAAAATATTACGATAAATTAGTTTTAAAAGT
GTTGCTCATAAATTGAAATCGAATGAGTCCTATAATTGTAGCATTGCAAACGACGAATCC
CAATATTAAACCTACACCTTAAAAACGAAGGATATGCGGCCGCGTAGTGC
gBlock_Synj_LHA:
CGGCCGCTATAACTAGTAACCATATTTAGGAAGCCATTGATGTCCTGCTAGTGGGCTCCA
CGCTTAGCTCGGAGCTTGCGGATCGGGCTCGCATCCTACTGCCCTCCAATATGTTGCATGC
ACCTACCACTGTGTTGAGAGAGCTATGCAAGCGCTACACTGAATATGTGCGTCCTCGAAT
GGCACGTGTAGCCGTGGGTACCTATAACGTCAACGGCGGCAAGCACTTCCGCAGCATTGT
GTTCAAGGATTCGCTGGCCGATTGGCTGCTCGACTGCCATGCCCTTGCCCGCTCCAAGGC
GCTTGTAGATGTGAACAATCCGTCGGAGAACGTCGATCATCCGGTGGATATCTACGCCAT
TGGATTCGAGGAGATTGTGGATCTGAATGCTTCCAACATAATGGCGGCCAGCACCGACAA
TGCCAAGTTGTGGGCCGAGGAGCTGCAGAAAACGATCTCGCGGGACAATGACTACGTGC
TGCTCACATACCAGCAACTGGTGGGCGTGTGCCTATACATCTACATCCGACCGGAGCACG
CGCCGCACATCCGGGACGTGGCCATCGACTGTGTTAAGACAGGATTGGGTGGTGCCACTG
GGAATAAGGGTGCCTGTGCCATTCGATTTGTGCTTCATGGTACTTCCATGTGCTTCGTGTG
TGCCCACTTTGCAGCCGGACAGTCACAGGTGGCTGAAAGGAACGCTGACTACGCGGAAA
TCACCCGGAAGCTGGCCTTCCCGATGGGCAGGACGCTAAAATCACACGACTGGGTGTTTT
GGTGCGGCGACTTCAACTATCGCATCGACATGGAGAAGGACGAATTAAAGGAGTGCGTA
CGTAATGGAGATCTCTCAACCGTCCTCGAGTTCGATCAATTGCGCAAGGAGCAGGAGGCT

GGCAATGTGTTTGGCGAATTCCTCGAGGGAGAGATCACTTTCGACCCGACGTACAAGTAT GATTTGTTCAGCGACGACTAGTAGGGATAACAGGGTAATG

***Gene: VPS35, Construct: pWhite-STAR_VPS35***
LHA_F_Primer: AACAGGGTAATGGTACCTACCACGCAATATCCACCACA
LHA_R_Primer: GCACTACGCGGCCGCATATCCTGTTTCTTAAGTTTTTGAACGA
RHA_F_Primer: CGGCCGCTATAACTAGTAACAAAATTAAAGCGTCTTTATACAAC
RHA_R_Primer: CATTACCCTGTTATCCCTACCTGAGCCAAGCTCAGCAG

***Gene: VAC14, Construct: pWhite-STAR2_VAC14***
LHA_F_Primer: GTCTCTAATTGAATTAGATCCCCCGTGGAGCAGGCGCCCGC
LHA_R_Primer: GCGGCCGCTATAACTAGTAACTGCTACTCACTTCTCGATC
RHA_F_Primer:
GGCACTACGCGGCCGCATATCTATTTGTGACCATTAGTCATCAAAATCGTGTTCCCATTTC AA
RHA_R_Primer: TAACAGGGTAATGGTACCTACCTGTGAAAGAACGTCCATA

***Primers used to verify the correct integration of the IMCE knock-out cassette***

Knock-ins made with pWhite-STAR used the combination of the F_LHA_junc_primer with R_w_start_primer and R_RHA_junc_primer with F_w_end_primer for verifying a correct cassette integration.

Knock-ins made with pWhite-STAR(II) used the combination of the F_LHA_junc_primer with F_w_end_primer and R_RHA_junc_primer with R_w_start_primer for verifying a correct cassette integration. This distinction is due to the reversed orientation of the mini_white cassette in the pWhite-STAR(II) construct.

F_w_end_primer: TTCGGAGTGATTAGCGTT
R_w_start_primer: GAGTGAGAGGTAATCGAA

| ***Gene*** | ***F_LHA_junc_primer*** | ***R_RHA_junc_primer*** |
|---|---|---|
| *ATP13A2* | GAGAGCAAAGGAAAAGGAA | CTAGTCTTTAGCCGCTTTT |
| *Vham8.9* | TGTATATGTCTGGTGTAGG | TTGTAGGGATTGGTGGTG |
| *CHCHD2* | GAAACATCGTCCCTTTTGTA | TCTTGTGGAGGATGTGGT |
| *Coq2* | AGTCGGACGAGATATTTG | GAGCGCGAGGAATGGAAA |
| *Dj1-a* | CAGAAAGGAATCGCATTG | CTGTTGCTTTGGTTGTTG |
| *Dj1-b* | GAAAGAGAGGGAAGACAGAA | CCCACTACAACTACAACA |
| *rme-8* | CCGTCCGTCTTTATCTTTA | GTCGTTGGTGCTGTAGTT |
| *auxilin* | CACTGCACCCGGAAAATCA | GGTAGCACAGGAAGTAGAG |
| *eIF4G1* | GAATTGAACGCTGGCATA | TTCCCCATTTTTTTCGCC |
| *ntc* | TTTAGTTACATTAGGGCTGG | GGGAAACCGAAAACATAG |
| *Gba1a_1b* | GCGTTATTTTGCCCACAT | ACTTGAGGTGATAGAGGG |
| *punch* | GTGGGTTTCAAGGATATG | TGTGAGTGCTAAAGTGGT |
| *Gdh* | TTTCGTTTGGTTCCCTGT | TTAAGGATGGATAGGTGG |
| *Omi* | GAAGAAACACGATTTGGGA | AGACAGACGGATGGTAGG |
| *Lrrk* | TCGATATAGGTTACAGTTGG | TATTCGCGTGATAGCTGT |
| *Tango14* | GTGCAATAGCCAAGCTTA | AAGTCAGGTGAGTGTAGAA |
| *Park* | AACACAAGCATACCGGAA | TTGTGCTGACTTTGATGG |
| *Pink1* | GGTCTGGGCGAACTAATG | CACCTCGTCGCAAAAGAA |
| *iPLA2-VIA* | CTGCTGGACAATGGGTAT | TTTACATACTGTGCCGGTT |
| *Loqs* | GAGAAACAGGGCTTTAAG | GCCCGTTGCATTTTTTAG |
| *Rab39* | TATAGGAAAACACTCGGG | GGAGTACTGGTGGGTCATA |
| *Synj* | CATATAGAATGCTGTCGT | CTGTTTCTCCGAGGTGTC |

| | | |
|---|---|---|
| *VPS35* | AAGGCGTTGTTGTAGAAG | AAGGCGTTGTTGTAGAAG |
| *VAC14* | TGTGCCCCAGATAGAAGT | TGTGCCCCAGATAGAAGT |
